# Tick extracellular vesicles alter keratinocyte function in the skin epidermis

**DOI:** 10.1101/2023.11.10.566612

**Authors:** Liron Marnin, Luisa M. Valencia, Haikel N. Bogale, Hanna J. Laukaitis-Yousey, Agustin Rolandelli, Camila Rodrigues Ferraz, Anya J. O’Neal, Axel D. Schmitter-Sánchez, Emily Bencosme Cuevas, Thu-Thuy Nguyen, Brenda Leal-Galvan, David M. Rickert, M. Tays Mendes, Sourabh Samaddar, L. Rainer Butler, Nisha Singh, Francy E. Cabrera Paz, Alejandra Wu-Chuang, Jonathan D. Oliver, Julie M Jameson, Ulrike G. Munderloh, Adela S. Oliva Chávez, Albert Mulenga, Sangbum Park, David Serre, Joao H.F. Pedra

## Abstract

Wound healing has been extensively studied through the lens of inflammatory disorders and cancer, but limited attention has been given to hematophagy and arthropod-borne diseases. Hematophagous ectoparasites, including ticks, subvert the wound healing response to maintain prolonged attachment and facilitate blood-feeding. Here, we unveil a strategy by which extracellular vesicles (EVs) ensure blood-feeding and arthropod survival in three medically relevant tick species. Through single cell RNA sequencing and murine genetics, we demonstrate that wildtype animals infested with EV-deficient *Ixodes scapularis* display a unique epidermal sub-population with a mesenchymal-like transcriptional program and an overrepresentation of pathways connected to wound healing. Furthermore, tick EVs inhibit proliferation and diminish the capacity of wound closure in keratinocytes. This occurrence was linked to phosphoinositide 3-kinase activity, keratinocyte growth factor 1 (KGF-1) and transforming growth factor β (TGF-β) levels. Collectively, we uncovered a strategy employed by a blood-feeding arthropod that disrupts the circuitry in cutaneous wound healing, contributing to ectoparasite fitness.

## INTRODUCTION

Ectoparasitic arthropods obtain their nourishment by feeding on a vertebrate host, providing an avenue for microbial transmission during hematophagy (Sonenshine and Roe, 2014). In North America, tick encounters account for approximately 77% of all arthropod-borne diseases, with most interactions attributed to *Ixodes*, *Amblyomma* and *Dermacentor* species (Eisen, 2022, Rosenberg et al., 2018). Notably, *Ixodes scapularis* is an arthropod vector of several human illnesses, including Lyme disease (Eisen, 2022). The Lone star tick *Amblyomma americanum* transmits bacteria that cause ehrlichiosis, while the American dog tick *Dermacentor variabilis* and *Dermacentor andersoni* carry pathogens associated with Rocky Mountain spotted fever and tularemia (Eisen, 2022, Rosenberg et al., 2018).

The unique microenvironment generated at the skin interface during a tick bite is conducive to arthropod hematophagy and pathogen transmission (Wikel, 2013). Unlike other blood-feeding arthropods, the development of hard ticks incorporates a series of events which necessitate long-term attachment to the skin (Sonenshine and Roe, 2014). This prolonged disruption of the host physical barrier poses a new challenge for tick survival as defense mechanisms are engaged. Tick saliva has been shown to be critical to subvert inflammation, blood coagulation, and nociception and antagonizes host immunity to enable attachment to the skin (Esteves et al., 2017, Francischetti et al., 2009, Kazimirova and Stibraniova, 2013, Kotal et al., 2015, Kotsyfakis et al., 2007, Kramer et al., 2011, Poole et al., 2013, Ribeiro et al., 1992, Ribeiro et al., 1985, Ribeiro et al., 1988, Simo et al., 2017, Valenzuela et al., 2002). Observations across various species further demonstrate a conserved ability whereby tick effectors perturb pro-inflammatory mediators responding to ectoparasite feeding (Bakshi et al., 2019, Dickinson et al., 1976, Esteves et al., 2017, Karim and Ribeiro, 2015). Such antagonism of host responses has been implicated in the dissemination and persistence of vector-borne pathogens (Chen et al., 2014, Kotsyfakis et al., 2010, Oliva Chávez et al., 2021).

As the largest organ in the body, the skin serves as the first line of defense against arthropod infestation and pathogen transmission (Eyerich et al., 2018, Glatz et al., 2017, Kabashima et al., 2019). The skin is comprised of three primary layers: the outermost epidermis, the underlying dermis, and the hypodermis or subcutaneous fat (Eyerich et al., 2018). The complex architecture and specialized cell populations comprising the skin affords the mammalian host a protective barrier against environmental and microbial threats, in addition to aiding in thermoregulation and prevention of trans-epidermal water loss (Kabashima et al., 2019, Proksch et al., 2008). During injury, such as laceration of the skin by a tick hypostome, various immune and sentinel cells are activated and release soluble factors that prompts the highly complex and intricate process of wound healing (Singer and Clark, 1999). Proper wound healing requires a high degree of coordination to orchestrate a response to an insult, which is broadly comprised of four overlapping stages: hemostasis, inflammation, proliferation, and tissue remodeling (Peña and Martin, 2024).

Little attention has been paid to the latter phases of skin healing during tick hematophagy, including proliferation, which promotes the process of re-epithelialization. Re-epithelialization, which is largely dependent on the migration and proliferation of keratinocytes, culminates in the regeneration of the epidermal-dermal junction and restoration of barrier integrity (Pastar et al., 2014, Rousselle et al., 2019). Keratinocytes function as structural cells which serve as the outermost layer in mammals (Pastar et al., 2014, Rousselle et al., 2019). Keratinocytes have also been recognized as sentinels, facilitating crosstalk with immune cells, and partaking in the initiation of the wound healing response upon injury (Piipponen et al., 2020). Dysfunction in keratinocyte-mediated closure has been reported in chronic wounds indicating their crucial role in skin homeostasis (Pastar et al., 2014, Wikramanayake et al., 2014).

The current paradigm at the tick-skin interface is that salivary molecules are deposited within the dermis during feeding, where they actively regulate the cutaneous response to an insult (Bernard et al., 2020, Wikel, 2013). The impact of tick feeding on the epidermis, which interfaces with the external environment, has been mostly neglected. The significance of the epidermis in countering tick infestation was documented in the late 1970s wherein Langerhans cells were shown to respond to salivary antigens (Allen et al., 1979). We also implicated extracellular vesicles (EVs) originating from the tick *I. scapularis* in promoting tick fitness and generating distinct outcomes of pathogen transmission in mammals. This mechanism was accomplished through the tick SNARE protein vesicle associated membrane protein 33 (Vamp33) and epidermal γδ T cells (Oliva Chávez et al., 2021).

In this article, we combined single cell RNA sequencing (scRNA-seq), murine genetics, intravital microscopy and flow cytometry to reveal that tick EVs disrupt principal components of the cutaneous wound healing circuitry. We discovered a unique epidermal sub-population in wildtype animals with a mesenchymal-like transcriptional program and an overrepresentation of pathways connected to wound healing during a bite from EV-deficient ticks. We further underpinned this biological network by demonstrating that tick EVs disrupt proliferation in keratinocytes and affect the levels of keratinocyte growth factor 1 (KGF-1), also named fibroblast growth factor 7 (FGF-7), and transforming growth factor β (TGF-β). Collectively, we illustrate a tick-induced interference of wound healing parameters via the skin epidermis, contributing to the process of arthropod hematophagy.

## RESULTS

### Tick extracellular vesicles enable arthropod fitness

We previously observed that EVs derived from *I. scapularis* enabled hematophagy (Butler et al., 2024, Oliva Chávez et al., 2021). We sought to corroborate our findings in other tick species of public health importance and assess the impact of EVs on arthropod fitness. We employed our validated methodology for the disruption of EV biosynthesis in ticks by silencing the expression of *vamp33* through RNA interference (RNAi) (Oliva Chávez et al., 2021) (Supplementary Table S1). Total genetic ablation in ticks remains beyond current technical capabilities because editing through clustered regularly interspaced short palindromic repeats (CRISPR) has only been applied to score morphological phenotypes, but not signaling pathways (Sharma et al., 2022).

We designated arthropods that had reduced *vamp33* gene expression as si*V33*, EV-deficient ticks, and the scramble control treatment as sc*V3*3, EV-sufficient ticks. Si*V33* and sc*V33* microinjected nymphs were placed on C57BL/6 mice and allowed to feed for 3 days (Figure 1a). On day 3, *I. scapularis* were assessed for efficiency of *vamp33* silencing, attachment, and collected for weight and post-feeding survival (Figure 1b-e). We observed a statistically significant difference in attachment between si*V33* and sc*V33* nymphs (Figure 1c) compared to our previous evaluation (Oliva Chávez et al., 2021). Diminished feeding was also measured for EV-deficient ticks as demonstrated by a 53% reduction in tick weight (Figure 1d). Interrupted feeding in *I. scapularis* led to reduced survival post-detachment (Figure 1e). An EV-associated fitness cost upon *vamp33* silencing was observed in all three clinically relevant tick species (Figure 1b-m). A notable exception was the lack of phenotypic differences in attachment for *A. americanum* compared to *I. scapularis* and *D. variabilis* (Figure 1c, g, and k). Collectively, these findings offer the prospect of a cross-species integrated management for mammalian infestation despite the distinct tick phylogeny.

**Figure 1:**
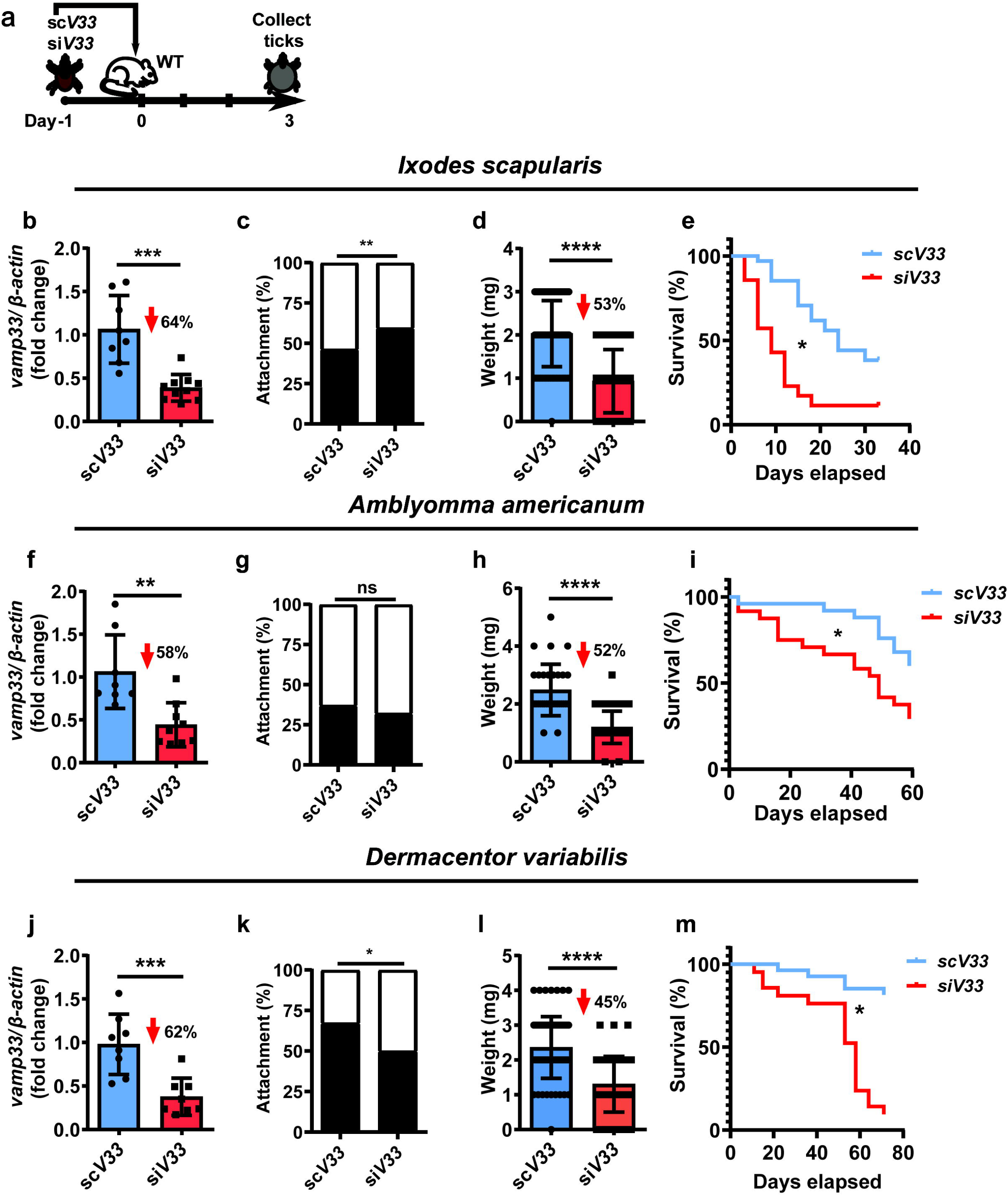
Tick EVs affect hematophagy and survival. **(a)** Graphical illustration of experimental design. **(b-m)** *Vamp33* siRNA (*siV33*) (red) or *vamp33* scramble control (*scV33*) (blue) microinjected nymphs were placed on C57BL/6 mice and allowed to feed for 3 days. On day 3, ticks were harvested and assessed for fitness measurements. Efficiency of *Vamp33* silencing, tick attachment, weight, and survival curves for **(b-e)** *I. scapularis,* **(f-i)** *A. americanum* and **(j-m)** *D. variabilis.* Graphs represent at least three independent experiments combined. Statistical significance shown as \**p*<0.05, \*\**p*<0.01, *** *p*<0.001 \*\*\*\**p*<0.0001. **(b, f, j, d, h, l)** Mann Whitney test with results presented as the mean +/- SD; **(c, g, k)** Fisher’s exact test and **(e, i, m)** Log-rank (Mantel-Cox) test. ns = not significant.

### Tick EVs alter epidermal immune surveillance

Recently, we reported that tick EVs within saliva affect the frequency of dendritic epidermal T cells (DETC) and alter the cytokine and chemokine milieu of the skin (Oliva Chávez et al., 2021). DETC surveillance of keratinocytes via various cell surface receptors is critical in a wounding response, leading to the activation and recruitment of immune cells, stimulation of keratinocytes for proliferation and survival, and anti-microbial responses (Jameson et al., 2002, Jameson et al., 2004, Keyes et al., 2016, Macleod and Havran, 2011, Sharp et al., 2005) (Figure 2a). This crosstalk and surveillance between DETC and keratinocytes led us to reason that tick EVs might not solely impact DETCs, but also likely influence the most abundant epidermal cell, the keratinocyte. Hence, we allowed EV-deficient (si*V33*) and EV-sufficient (sc*V33*) *I. scapularis* nymphs to feed on mice for 3 days and collected the skin biopsy for flow cytometry evaluation (Supplementary Figure S1). We observed a decrease in DETC frequency during sc*V33* tick feeding on mice compared to naïve skin (Figure 2b). Conversely, DETC frequency remained at homeostatic levels after impairment of tick EVs (si*V33*) and ectoparasite feeding on murine animals (Figure 2b).

**Figure 2:**
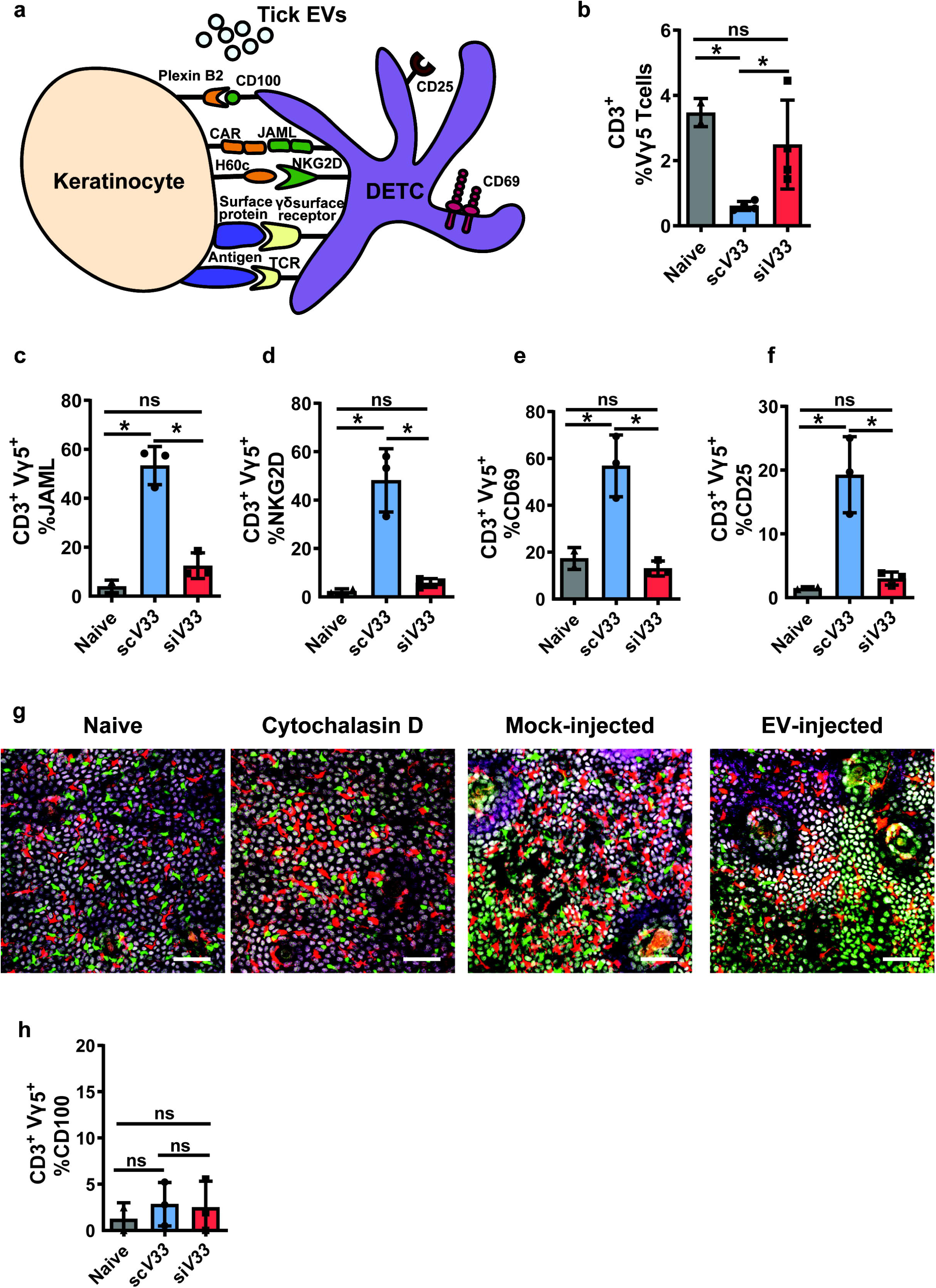
Tick EVs alter epidermal immune surveillance. **(a)** Schematic representation of the DETC-keratinocyte crosstalk at the skin epidermis. **(b-f, h)** *I. scapularis scV33* (blue) or *siV33* (red) ticks were placed on C57BL/6 mice and allowed to feed for 3 days. On day 3, biopsies were taken from the skin at the bite site and compared to the naïve treatment (gray). **(b)** DETC (Vγ5), **(c)** JAML, **(d)** NKG2D, **(e)** CD69, **(f)** CD25, and **(h)** CD100 cells were assessed by flow cytometry. Graphs represent 1 of 3 independent experiments. **(g)** Epidermis containing Langerhans cells (red), DETCs (green), and keratinocytes (white) imaged on day 3 after injection with phosphate buffered saline (PBS - mock) or EV (4x10^7^ particles) into the mouse ear. Cytochalasin D (100 µg) was applied topically on the mouse ear every 24 hours for 2 days to induce DETC rounding as a positive control. Langerhans cells, DETCs and epithelial cells were simultaneously visualized in the *huLangerin-CreER; Rosa-stop-tdTomato; CX3CR1-GFP^+/−^; K14-H2B-Cerulean* mouse strain. Cre expression was induced with an intraperitoneal injection of tamoxifen (2 mg). The size of the scale bar represents 50 μm. Images from one out of three independent experiments. Statistical significance shown as \**p*<0.05, ns = not significant. Data are presented as a mean +/- SD. Significance was measured by One-way ANOVA followed by Tukey’s *post hoc* test.

DETCs exhibit a dendritic shape that allows for continuous surveillance of neighboring keratinocytes through various receptor-ligand interactions (Jameson et al., 2002, Witherden et al., 2012) (Figure 2a). Upon tissue damage, stressed keratinocytes upregulate ligands and antigens that stimulate DETCs in a non-major histocompatibility complex (MHC)-restricted manner (Havran et al., 1991). Activated DETCs will then alter their morphology by retracting dendrites and assuming a rounded configuration to facilitate migration to the site of injury (Jameson et al., 2002, Nielsen et al., 2017). To determine the possible role of keratinocytes during tick feeding, we assessed the DETC co-stimulatory markers that facilitate immune surveillance. We observed an elevated co-receptor frequency among DETCs found at the skin interface where EV-sufficient ticks fed on mice, including the junctional adhesion molecule-like (JAML) and the C-type lectin-receptor NKG2D (also known as KLRK1) (Girardi et al., 2001, Whang et al., 2009) (Figure 2c-d). Similar findings were also observed for the activation markers CD69 and CD25 (Figure 2e-f). Conversely, JAML, NKG2D, CD69 and CD25 were not upregulated in the bite of EV-deficient ticks during murine feeding (Figure 2c-f).

Morphologically, the hallmark of DETC activation is the conversion of a dendritic to a rounded morphology that facilitates intraepidermal migration, a phenomenon that is partially regulated by CD100 signaling (Thelen and Witherden, 2020, Witherden et al., 2012). To capture morphological changes in DETCs, we employed intravital microscopy of EV injection into the ear of a triple-reporter mouse model. Intravital microscopy of EV injection into the ear of this mouse model revealed that tick EVs did not promote rounding of DETCs, as compared to the positive control cytochalasin D (Figure 2g). Supporting epidermal intravital imaging findings, expression of CD100 was not altered during a tick bite regardless of the EV status (Figure 2h). Altogether, these findings provided evidence that tick EVs functionally alter immune surveillance of the epidermal niche by DETCs.

### ScRNA-seq characterization of epidermal cells during tick feeding

Given the functional perturbations in DETCs during tick feeding, and the well documented importance of the DETC-keratinocyte crosstalk during wounding, we hypothesized that the epidermal healing circuitry is likely being altered during tick feeding. To evaluate this hypothesis, we utilized scRNA-seq to analyze the impact of tick EVs on the epidermal immune environment in both DETC-deficient (FVB-Tac) and DETC-sufficient (FVB-Jax) mice three days after tick feeding.

FVB-Tac mice are depleted of functional DETCs due to a failure of thymic selection because of a natural mutation of the *skint1* gene, consequently obstructing DETC seeding of the epidermis (Barbee et al., 2011, Boyden et al., 2008, Lewis et al., 2006). Importantly, FVB-Tac mice is an acceptable DETC-deficient mouse model by the scientific community (McKenzie et al., 2022, Munoz-Ruiz et al., 2023) and enables the selective study of Vγ5Vδ1 epidermal γδ T cells, while limiting non-specific results potentiated by dermal γδ T cells. Skin punch biopsies were obtained from the bite site, and the epidermis was enzymatically separated from the dermis. Live cells were sorted by fluorescence activation and libraries were generated for Illumina sequencing (Figure 3a).

**Figure 3:**
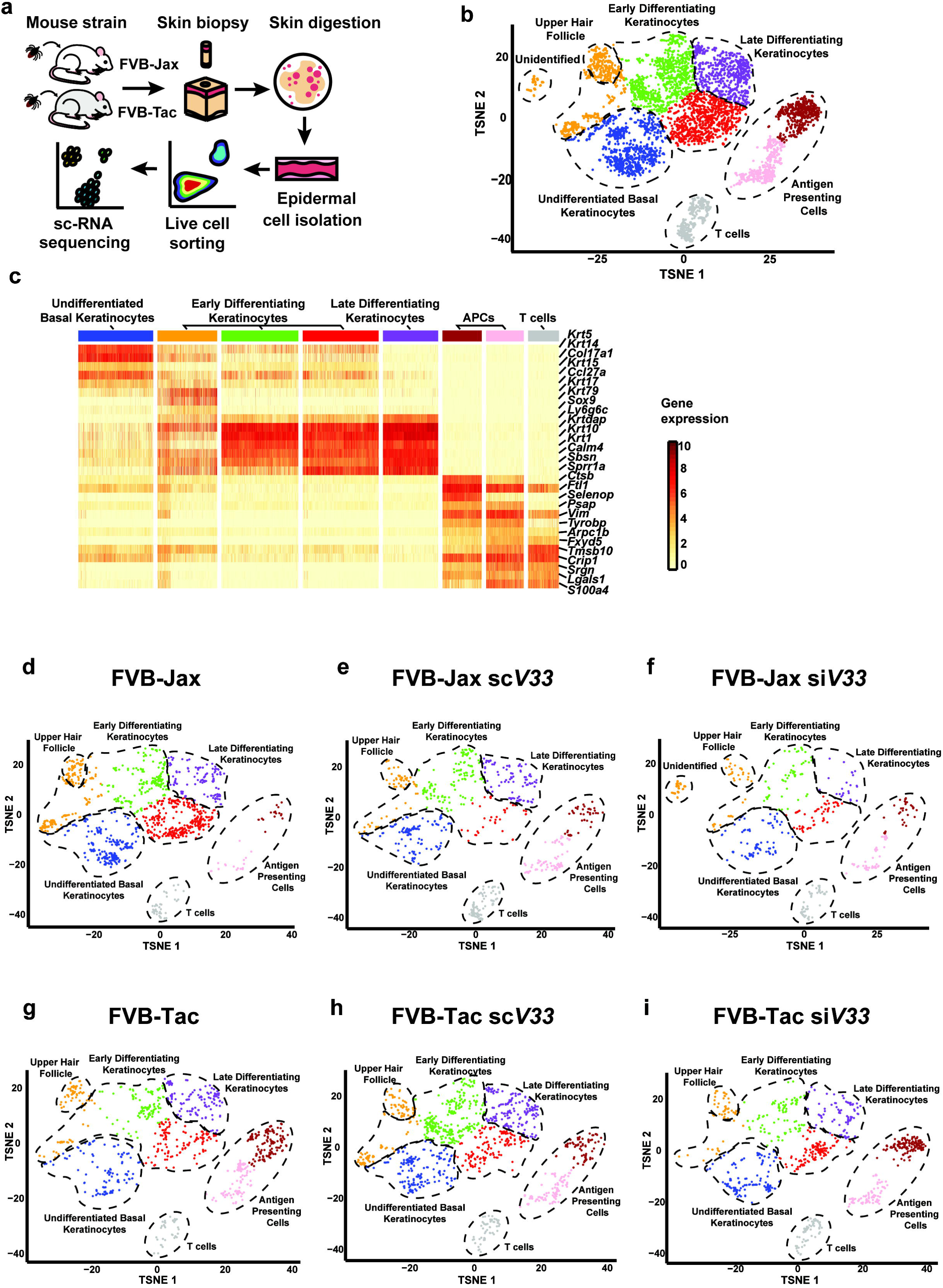
Epidermally-enriched scRNA-seq of the tick bite site. **(a)** Overview of the experimental design. *ScV33* and *siV33 I. scapularis* nymphs were placed on FVB-Jackson (FVB-Jax) or FVB-Taconic (FVB-Tac) mice and fed for 3 days. Skin biopsies at the bite site were digested with dispase and collagenase for epidermal cell isolation. Cells were sorted and prepared for scRNA-seq. **(b)** Composite tSNE plot of keratinocyte, T cell and antigen presenting cells in FVB-Jax and FVB-Tac mice in the presence or absence of *I. scapularis* nymphs microinjected with *scV33* or *siV33*. tSNE plot represents 5,172 total cells following filtration as described in the materials and methods. **(c)** Heatmap depicting expression of the top 5 marker genes present in clusters from the epidermally enriched tSNE plot clusters (as shown in b). **(d-i)** Individual tSNE plots separated by mouse strain (FVB-Jax or FVB-Tac) in the presence or absence of *I. scapularis* nymphs microinjected with *scV33* or *siV33*.

Our analysis encompassed approximately 20,640 cells, with an average of 88,027 reads. Our initial investigation resulted in 23 clusters (Supplementary Table S2). Next, we applied a fixed threshold to retain cells with more than 2500 UMIs (Supplementary Figure S2a-b) and applied the DoubletFinder R package to predict doublets (Supplementary Figure S2c-d). We identified 10 distinct groups of cells through an analysis of marker genes within each cluster relative to the entire dataset (Supplementary Table S3). Keratinocytes, T cells, fibroblasts and endothelial cells were observed in our scRNA-seq results (Supplementary Figure S2d). The presence of dermal clusters in our study was likely due to an incomplete epidermal-dermal border separation during the enzymatic dissociation of skin biopsies. Thus, we subjected keratinocytes, T cells, and antigen-presenting cells (APCs) to a second round of clustering (Supplementary Table S4). This dataset revealed a total of 8 clusters visualized in t-distributed stochastic neighbor embedding (t-SNE) (Figure 3b) for a total of 5,172 total cells with a median UMI count of 13,910 per cell.

Throughout the process of differentiation, keratinocytes express different types of keratins, including keratins (*Krt*) *1*, *5*, *10*, and *14* (Fuchs, 1993). Elevated levels of *Krt5* and *Krt14* expression enabled the recognition of undifferentiated cells residing within the basal layer of the epidermis (Figure 3c, Supplementary Table S5). *Krt1*, *Krt10*, and *Involucrin* were used to discern early and late-stage differentiation of keratinocytes (Figure 3c, Supplementary Figure S3, Supplementary Table S5). APCs and T cells were identified by the T cell receptor alpha constant (*Trac*), the T cell receptor delta constant (*Trdc*), and the histocompatibility class II antigen (*H2-Aa*) (Supplementary Table S4 and Supplementary Table S6). The mouse epidermis harbors hair follicles with distinct physiological functions (Joost et al., 2018, Joost et al., 2016). Our dataset only accounted for compartments in anatomical proximity to the epidermis (Supplementary Figure S4, Supplementary Table S6).

We then determined the percent distribution of interfollicular epidermal cells per treatment. In the skin biopsy where ticks fed on immune intact mice (FVB-Jax sc*V33* and FVB Jax si*V33*), we observed a decrease in keratinocytes and an overrepresentation of T cells and APCs compared to the naïve skin (Supplementary Figure S5a, Supplementary Table S7). A similar effect was not observed when ticks fed on the skin of DETC-deficient mice (Supplementary Figure S5b, Supplementary Table S7), presumably due to the diminished wound healing capacity observed in FVB-Tac animals (Keyes et al., 2016). We confirmed the depletion of DETCs in the epidermis of FVB-Tac mice. Gene expression of *Trdv4* in the T cell cluster, which encodes for the receptor Vδ1 in DETCs, was reduced in FVB-Tac compared to the FVB-Jax mouse strain (Supplementary Figure S5c). Notably, partitioning of epidermal clusters by experimental conditions revealed an unidentified epidermal subcluster found solely when EV-deficient ticks fed on FVB-Jax mice (Figure 3d-i). The presence of this distinct cluster raised the hypothesis that EVs might exert an influence on keratinocytes within the context of DETCs, given its absence in FVB-Tac mice.

### Tick EVs impact an epidermal sub-population with a prominent wound healing signature

The emergence of this unique epidermal subcluster responding to si*V33* tick feeding prompted us to further investigate their role by subjecting all keratinocytes to a subsequent round of clustering. This examination revealed keratinocyte populations at various differentiated states and highlighted the presence of an unidentified epidermal population exclusive to EV-deficient tick feeding on FVB-Jax mice (Figure 4a, Supplementary Figure S6, Supplementary Table S8).

**Figure 4:**
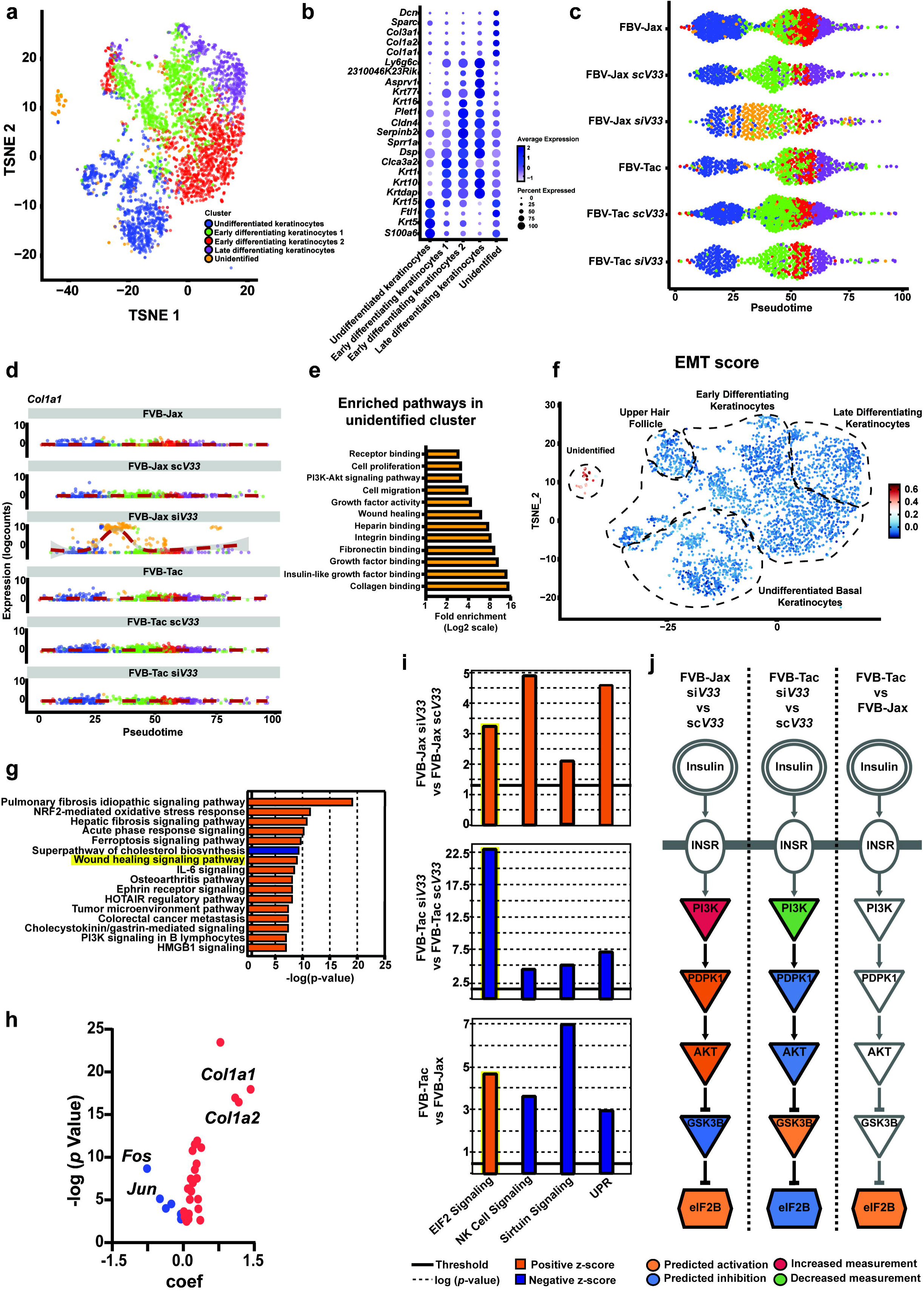
Tick EVs impacts the keratinocyte transcriptional program. **(a)** Composite tSNE plot of keratinocytes in FVB-Jax and FVB-Tac mice in the presence or absence of *I. scapularis* nymphs microinjected with *scV33* or *siV33*. **(b)** Dot plot of the top 5 marker genes present in the keratinocyte clusters (as shown in a). Average gene expression is demarked by the intensity of color. Percent of gene expression within individual clusters is represented by the dot diameter. **(c)** Cells colored by clusters originated from the keratinocyte tSNE plot (as shown in a) ordered across pseudotime (x-axis) for naïve, *scV33*-, and *siV33*-tick bites of FVB-Jax and FVB-Tac mice. **(d)** Expression of *Col1a1* across treatments ordered across pseudotime (x-axis) for naïve, *scV33*-, and *siV33*-tick bites of FVB-Jax and FVB-Tac mice. **(e)** Enriched pathways in the unidentified cell cluster based on functional annotation in DAVID. Fold enrichment is indicated in a Log2 scale. \**p* value and false discovery rate (FDR)<0.05 were set as threshold. KEGG, GO and InterPro were used as reference annotation databases. **(f)** The “AddModuleScore” function was used to generate an epithelial-to-mesenchymal transition (EMT) score across all keratinocytes based on 102 epithelial-to-mesenchymal transition genes. Selected genes were reported as indicative of keratinocytes in intermediary or mesenchymal transitioning states. **(g)** Ingenuity pathway analysis comparing keratinocytes of skin biopsies from FVB-Jax *siV33* to FVB-Jax *scV33*. Blue denotes pathways predicted to be inhibited (negative z-score) whereas orange indicates pathways predicted to be activated (positive z-score) based on default parameters. Differential expression datasets were assessed for canonical pathway analysis. Results are shown in a -log (*p*-value) scale. \**p* value and FDR< 0.05 were set as threshold. **(h)** Volcano plot of genes representing the wound healing signaling pathway in keratinocytes of FVB-Jax *siV33* compared to FVB-Jax *scV33* datasets (highlighted in yellow; g). Blue denotes decrease whereas red indicates increase in the coefficient (coef) of expression. **(i)** Ingenuity pathway analysis derived from *siV33* compared to the bite of *scV33* ticks on FVB-Jax or FVB-Tac mice. Canonical pathways predicted to be inhibited (blue, negative z-score) or activated (orange, positive z-score) based on differential expression profile. The solid line indicates the *p* value significance threshold of 0.05 (-log=1.3). **(j)** The signaling cascade of EIF2 (highlighted in yellow, i), yielding (→) or inhibitory (┤) arrows. Orange indicates activation whereas blue shows inhibition according to the IPA prediction. Gene expression based on the scRNA-seq experiment is indicated in red (increased) or green (decreased). Gray – denotes no expression or prediction.

Notably, this unidentified epidermal subcluster arising during EV-deficient tick feeding on FVB-Jax mice exhibited marker genes such as *Col1a1, Col1a2,* and *Col3a1* (Figure 4b).

The heterogeneity and plasticity of keratinocytes is crucial for various functions, both during homeostasis and in response to external stimuli (Dekoninck and Blanpain, 2019, Rice and Rompolas, 2020). Keratinocyte differentiation and response to wounding are dynamic biological processes in which gradual alteration in the transcriptional program has been explored to elucidate the coordination of broader cellular circuits (Joost et al., 2020, Joost et al., 2018, Joost et al., 2016, Reynolds et al., 2021). Thus, we employed pseudotime analysis to align keratinocytes along an inferred developmental trajectory based on their expression profile (Figure 4c). Gene expression signatures mirrored the sequence of differentiation, starting with markers associated with undifferentiated basal states to terminally differentiated keratinocytes (Supplementary Figure S7). The aforementioned epidermal subcluster was present along the pseudotime axis, proximal to the basal keratinocyte cluster, and was also exclusive to the condition where EV deficient ticks fed on FVB-Jax mice (Figure 4c). Evaluation of the marker gene *Col1a1* across pseudotime further underscored the distinct transcriptional program of this unidentified epidermal keratinocyte subcluster (Figure 4d). Altogether, these analyses reveal the transient state of this unidentified subcluster, which reflected a distinct expression profile that distinguished this subpopulation from other keratinocytes.

Given the distinct transcriptional profile of this unidentified subcluster, we sought to assess biological pathways enriched that may reveal the functional role of these cells. Pathway enrichment analysis of all significant marker genes in the unidentified subpopulation revealed gene signatures associated with both keratinocytes and fibroblasts, showcasing an overrepresentation of genes associated with the wound healing circuitry, including growth factor, collagen, fibronectin and heparin binding, and phosphoinositide 3-kinase (PI3K) activity (Figure 4e, Supplementary Table S9). These molecules have been implicated in keratinocyte function during wound healing, primarily by enhancing keratinocyte proliferation and migration to support re-epithelialization and tissue repair (Bártolo et al., 2022, Matsuura-Hachiya et al., 2018, Misiura et al., 2020).

Upon injury, keratinocytes adjacent to the wound are quiescent, opting for a migratory phenotype that allows for the initiation of re-epithelialization (Dekoninck and Blanpain, 2019). Conversely, keratinocytes farther from the wound edge undergo a proliferative burst, allowing for the closure of the gap generated by migratory keratinocytes (Aragona et al., 2017). To facilitate migration, basal keratinocytes at the leading wound edge undergo an epithelial-mesenchymal transition in which they adopt a mesenchymal-like transcriptional program and morphology (Chen et al., 2024, Haensel and Dai, 2018, Yao et al., 2024). The mesenchymal-like gene expression observed in this cluster and the transient state predicted by pseudotime analysis, prompted us to examine the possibility of this unidentified subcluster being keratinocytes undergoing epithelial-to-mesenchymal transition. As such, we assessed the distribution of 102 genes across our entire keratinocyte cluster to generate an epithelial-to-mesenchymal computational score (Supplementary Table S10) (Guo et al., 2024).

The epithelial-to-mesenchymal transition score distribution across the keratinocyte cluster revealed this unidentified subcluster to have a distinct enrichment for epithelial-to-mesenchymal transition genes compared to the remaining keratinocytes (Figure 4f). The mesenchymal-like transcriptional profile of keratinocytes undergoing epithelial-to-mesenchymal transition also has significant overlap with fibroblasts (Guo et al., 2024). On the one hand, assessing gene expression for epithelial-to-mesenchymal transition across cells within the complete unfiltered scRNAseq dataset revealed fibroblasts to have an elevated computational score due to their intrinsic mesenchymal identity. On the other hand, the unidentified subpopulation identified when EV deficient ticks fed on FVB-Jax mice showcased a diminished transcriptional signature compared to fibroblasts (Supplementary Figure S8). Collectively, our findings suggested that the unique epidermal subpopulation exclusive during EV-deficient tick feeding on FVB-Jax mice might be keratinocytes undergoing epithelial-to-mesenchymal transition. Unfortunately, limitations due to the rarity of this cutaneous subpopulation and the difficulty in obtaining epidermal tissue from a single tick-infested biopsy restricted any subsequent study.

To interpret the impact of feeding on murine skin between EV-deficient and EV-sufficient ticks, we subjected keratinocytes to a differential expression analysis and assessed enriched pathways through ingenuity pathway analysis (IPA). Notably, we observed a wound healing signature in the keratinocytes of DETC-sufficient mice infested with EV-deficient ticks, which were not detected in animals deficient for DETCs (Figure 4g). Further inspection of differentially expressed genes annotated for wound healing revealed a decrease in transcript levels for *Fos* and *Jun* and an increase of expression for *Col1a1* and *Col1a2* (Figure 4h, Supplementary Table S11). *Fos* and *Jun* are subunits of AP-1, which is important for epithelial proliferation, migration and differentiation (Angel et al., 2001, Gangnuss et al., 2004, Li et al., 2003). Conversely, collagen has various roles during all stages of wound healing, aiding in the regulation of the wound healing response, regeneration of a new basement membrane and extracellular matrix, reinforcing barrier integrity and facilitating the stratification of epidermal layers (Matsuura-Hachiya et al., 2018, Ohashi et al., 2025). Altogether, we provided evidence that tick EVs impaired wound healing through specific molecular pathways in keratinocytes.

To understand how tick EVs influenced wound healing parameters in keratinocytes, we evaluated molecular networks altered in the epidermis of DETC-sufficient and DETC-deficient mice. We performed a similar analysis in naive animals to exclude confounding effects originating from genetic differences occurring between these two strains. Four pathways were identified: eukaryotic Initiation Factor 2 (EIF2), natural killer (NK) cell, sirtuin signaling, and the unfolded protein response (UPR) (Figure 4i, Supplementary Table S12). The results obtained concerning NK cell, sirtuin signaling, and the UPR pathways were likely due to the *skint1* deficiency in FVB-Tac mice. However, the EIF2 cascade was dependent on tick EVs because the computational prediction occurred regardless of the mouse genetic background (yellow highlight, Figure 4i). A granular view of the EIF2 signaling pathway displayed PI3K as part of the biological circuit targeted by tick EVs (Figure 4j, Supplementary Table S13). The PI3K/Akt pathway is important for skin development and wound healing, contributing to keratinocyte proliferation, migration and differentiation (Calautti et al., 2005, Misiura et al., 2020, Teng et al., 2021). Collectively, our scRNA-seq analysis of the tick-skin interface suggested a disruption of the wound healing circuitry by tick EVs acting on keratinocytes.

### Tick EVs interfere with keratinocyte proliferation

The PI3K/Akt/mTOR pathway has been observed in the proliferative zone where keratinocytes are located and correlates with accelerated wound closure (Squarize et al., 2010, Teng et al., 2021). To evaluate whether tick EVs interfered with keratinocyte proliferation, we used the protein Ki-67 and flow cytometry as a readout for proliferative keratinocytes (Supplementary Figure S9). We observed a significant reduction in keratinocyte proliferation when EV-sufficient ticks fed on wildtype mice (Figure 5a). However, the effect of keratinocyte proliferation was not observed in the absence of tick EVs or DETCs, suggesting that the impact of tick EVs on keratinocyte proliferation was DETC dependent (Figure 5a).

**Figure 5:**
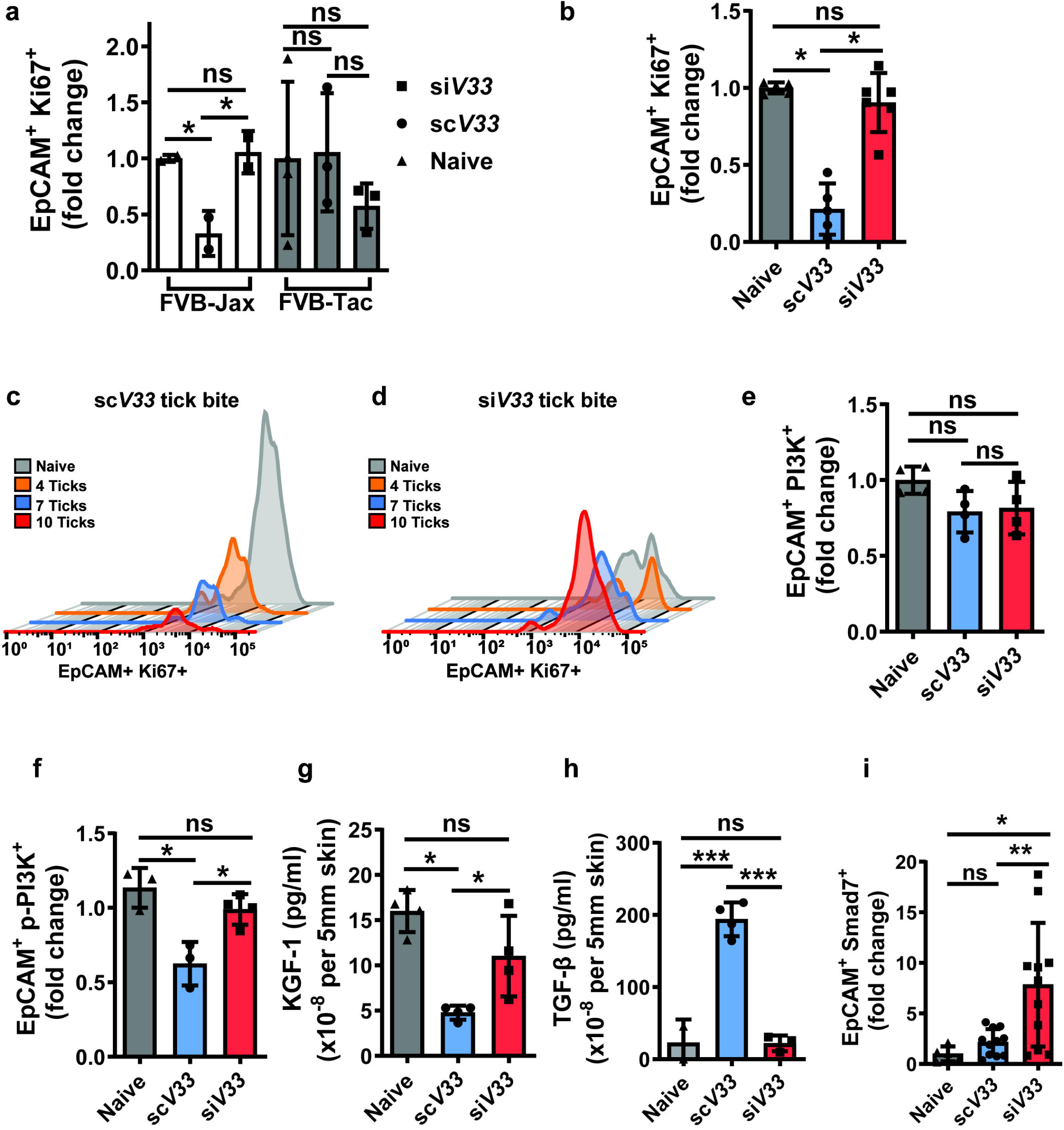
Tick EVs disrupt the cutaneous wound healing circuitry *in vivo*. **(a)** *ScV33* (circle) or *siV33* (square) injected *I. scapularis* nymphs were fed on FVB-Jax (white) or FVB-Tac (gray) mice for 3 days. Biopsies were taken from the skin at the bite site and assessed for EpCAM^+^ Ki67^+^ keratinocytes by flow cytometry. **(b-i)** *ScV33* or *siV33* ticks fed on C57BL/6 mice for 3 days. Biopsies were taken from the skin at the bite site and processed for flow cytometry and ELISA analysis. **(b)** EpCAM^+^ Ki67^+^ keratinocytes assessed by flow cytometry. Graph displays proliferation changes within the *scV33* or *siV33* treatments compared to the naïve skin. **(c-d)** Flow cytometry histogram plots of EpCAM^+^ Ki67^+^ keratinocytes. **(c)** *scV33* or **(d)** *siV33* treatments displayed according to the number of ticks bitten per biopsy. X-axis shows fluorescence intensity, and the Y-axis indicates the count of events in the fluorescence channel. **(e)** PI3K p85^+^, and **(f)** phospho-PI3K p85/p55^+^ keratinocytes were assessed by flow cytometry within the *scV33* or *siV33* treatments compared to the naïve skin. **(g-h)** ELISA analysis of **(g)** KGF-1 and **(h)** TGF-β levels normalized to total protein per 5 mm skin punch biopsy. **(i)** EpCAM^+^ Smad7^+^ keratinocytes assessed by flow cytometry. **(a-b, e-i)** Significance was measured by One-way ANOVA followed by Tukey’s post hoc test. All experiments have statistical significance shown as \*\*\**p*<0.001, \*\**p*<0.01, \**p*<0.05, ns = not significant. Data are presented as the mean +/- SD.

The genetic constitution of a mouse may lead to substantial alterations in phenotypic traits (Tanner and Lorenz, 2022, Woodworth et al., 2004). We therefore sought to corroborate our evidence of altered keratinocyte proliferation in C57BL/6 mice, which similarly revealed a significant reduction in the frequency of EpCAM^+^ Ki67^+^ keratinocytes during EV-sufficient tick feeding (Figure 5b). Remarkably, the ability of ticks to impair keratinocyte proliferation was observed in a density-dependent manner. As the number of ticks feeding on C57BL/6 mice increased, the capacity of keratinocytes to proliferate decreased (Figure 5c). This observation was not recorded in mice infested with ticks deficient for EVs (Figure 5d). To assess PI3K signaling in keratinocytes, we ascertained the keratinocyte PI3K status by flow cytometry due to its ability to assess protein expression on limited cell counts. Variation in the total PI3K in keratinocytes across treatments was not observed (Figure 5e). However, a decrease in phospho-PI3K-positive keratinocytes was recorded when ticks deficient in EVs fed on C57BL/6 mice (Figure 5f).

The tightly coordinated processes conducive to mammalian wound healing require a complex interplay of soluble factors that regulate the various wound healing parameters (Pastar et al., 2014). We therefore reasoned that the observed alterations in keratinocyte proliferation may be indicative of changes in soluble mediators necessitated for wound closure. Thus, we also assessed levels of KGF-1 and TGF-β, two growth factors that have a notable role regulating proliferation and epithelial-to-mesenchymal transition in keratinocytes (Pastar et al., 2014, Yao et al., 2024). Our findings revealed that the bite of *I. scapularis* ticks reduced KGF-1 and increased the level of TGF-β in the skin, compared to the EV-deficient treatment (Figure 5g-h).

Additionally, increased levels of TGF-β correlated to a significant decrease in the frequency of EpCAM^+^ keratinocytes expressing the negative regulator of TGF-β signaling, Smad7, in skin infested with *scV33* ticks compared to *siV33* ticks. (Figure 5i).

To further explore the regulatory role of ticks on keratinocytes, we then examined the consequences of EV stimulation on human keratinocytes in a scratch-wound assay. Briefly, HaCaT cell monolayers were subjected to a scratch-wound and stimulated with *I. scapularis* EVs derived from partially fed adult ticks as previously described (Oliva Chávez et al., 2021). The area of the scratch wound was monitored at 0, 12 and 24 hours to assess the percentage of gap closure based on the scratch area generated at 0 hours (Figure 6a). Stimulation with tick EVs resulted in a significant decrease in the closure gap at both 12 and 24 hours compared to the untreated culture medium (Figure 6b-c). Our *in vivo* findings suggested tick EVs to exert an inhibitory effect on keratinocyte proliferation. Thus, we assessed HaCaT cell proliferation at 24 hours post scratch and EV stimulation as a measurement of Ki67 protein expression using mitomycin C as a control through flow cytometry. Remarkably, stimulation with tick EVs significantly reduced Ki67 protein expression in human keratinocytes compared to the untreated culture medium (Figure 6d). Collectively, these results corroborate our mouse findings whereby tick EVs inhibit keratinocyte proliferation and regulate the epidermal wound healing circuitry, maintaining successful arthropod hematophagy.

**Figure 6:**
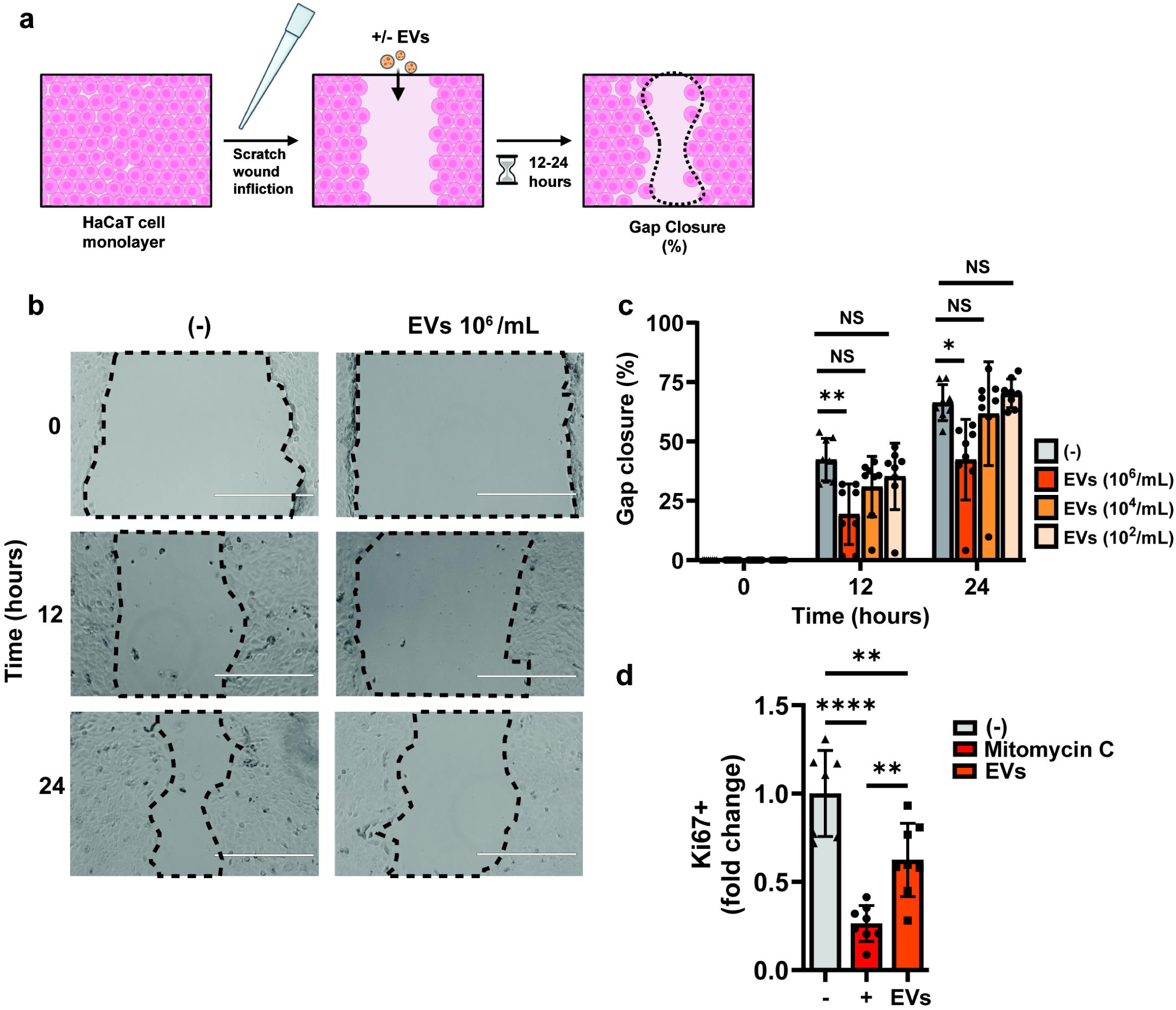
Tick EVs inhibit keratinocyte gap closure and proliferation. Human keratinocytes (HaCaT) monolayers were inflicted a scratch wound and stimulated with *I. scapularis* EVs. **(a)** Experimental design. **(b)** HaCaT monolayers were imaged at 0, 12 and 24 hours. Dashed line denotes scratch area, and images are representative of medium (-) or EV stimulated (10^6^/mL) cells. **(c)** Scratch area was quantified using ImageJ software and the percentage of gap closure at each time point was determined based on the original area of scratch inflicted at 0 hours for each treatment. The size of the scale bar represents 400 μm. Significance was measured by non-parametric Kruskal-Wallis test followed by Dunn’s multiple comparisons test. **(d)** Cells were treated with (-) medium, (+) mitomycin C or EVs. Ki67^+^ keratinocytes were assessed by flow cytometry 24 hours post-scratch infliction. Significance was measured by One-way ANOVA followed by Tukey’s post hoc test. All experiments have statistical significance shown as \*\*\*\**p*<0.0001, \*\**p*<0.01, \**p*<0.05. Data are presented as the mean +/- SD.

## DISCUSSION

Ticks are ancient hematophagous arthropods that co-evolved with their hosts for millions of years (Sonenshine and Roe, 2014). Currently, we have a limited understanding of the mechanisms employed by these ectoparasites to subvert wound healing in the skin during hematophagy. Previously, we implicated *I. scapularis* EVs in promoting tick fitness, affecting the frequency of DETCs in the skin, and generating distinct outcomes of pathogen transmission (Oliva Chávez et al., 2021). Here, we unveil that certain stages of the cutaneous wound healing response are a target of tick-mediated host immunomodulation.

By utilizing our previously established methodology for the knock-down of EV biosynthesis in ticks, we demonstrated the importance of EVs as a conserved strategy for arthropod fitness in three medically relevant tick species: *I. scapularis, A. americanum,* and *D. variabilis.* While current methodologies limit follow up studies in ticks, our previously published analysis characterizing EVs uncovered similarities and differences in cargo and post-translational modifications that were dependent on the source (*i.e.,* tick cells or salivary glands) (Oliva Chávez et al., 2021). These differences in the EV cargo might also be contingent upon the day of feeding, life stage and tick species. Upon future tool development, alterations in the cargo of tick EVs should be a source of scientific investigation in the context of how *I. scapularis* affect wound healing in the skin.

During skin injury, DETCs are activated whereby recognition of self-antigens on damaged or stressed keratinocytes through their TCR and co-stimulatory markers facilitate effector functions. (Jameson et al., 2004). Previous studies in animals devoid of DETCs unveiled delayed wound repair, highlighting their importance in orchestrating the cutaneous wound healing response (Jameson et al., 2004). Our work reveals that *I. scapularis* EVs led to a decrease in DETCs at the bite site. Furthermore, DETCs remaining in the epidermis displayed altered expression of co-stimulatory molecules. These findings suggested an altered DETC-mediated immunosurveillance in the skin during a tick bite.

The altered DETC co-stimulatory phenotype may be consequential to the capacity of keratinocytes to repair the wound (Jameson et al., 2002). To capture alterations in the cutaneous landscape during tick feeding, we employed a scRNAseq approach in FVB-Tac mice, an animal model wherein Vγ5Vδ1 γδ T cells are naturally absent in the skin epidermis (Barbee et al., 2011,

Boyden et al., 2008, Lewis et al., 2006). While we acknowledge that other animal models exist for the depletion of Vγ5δ1 γδ T cells, unfortunately, they also affect dermal Vγ6Vδ1 γδ T cells (Itohara et al., 1993, McKenzie et al., 2022, Sandrock et al., 2018).

Characterization of the epidermal response to tick infestation revealed a unique epidermal subcluster when EV-deficient ticks fed on DETC-sufficient mice. This epidermal subcluster exhibited an overrepresentation of pathways associated with collagen, fibronectin binding, and cell proliferation. Notably, pseudotime trajectory analysis revealed this subpopulation to be proximal to undifferentiated basal keratinocytes, with a distinct mesenchymal-like transcriptional program, which may reflect the transition adopted by migratory keratinocytes at the wound edge (Yao et al., 2024). The absence of this unique epithelial-to-mesenchymal transition population when EV-deficient ticks fed on DETC-deficient mice suggested a phenotype dependent of intraepithelial γδ T cells.

Computational analysis of the epidermal response to feeding of EV-deficient ticks on DETC-sufficient mice revealed specific biological signatures associated with the down regulation of AP-1 and upregulation of PI3K. These molecular circuits have been implicated in epithelial proliferation, migration and maintenance of barrier integrity (Angel et al., 2001, Gangnuss et al., 2004, Jochum et al., 2001, Li et al., 2003, Matsuura-Hachiya et al., 2018). The mammalian wound healing response is a tightly coordinated circuit requiring the temporal regulation of four stages: hemostasis, inflammation, proliferation and remodeling (Pastar et al., 2014). We showed the differential levels of TGF-β in keratinocytes during a tick bite. Whereas enhanced TGF-β levels in the skin bite of EV-sufficient ticks correlated to lower Smad7 levels, the skin where EV-deficient ticks were placed showcased KGF-1 and TGF-β levels comparable to naïve conditions. KGFs released by various cell types serve as a strong mitogenic factor for both mouse and human keratinocytes (Jameson et al., 2002, Werner et al., 1992). KGFs promote the process of re-epithelialization by stimulating keratinocyte migration and proliferation (Jameson et al., 2002), and its overexpression can lead to a hyperproliferative state associated with skin disorders (Ni and Lai, 2020).

TGF-β plays a critical role in wound healing, exerting a broad spectrum of functions to coordinate various stages of tissue repair (Liarte et al., 2020, Pastar et al., 2014). Increased levels of TGF-β in the epidermis have been associated with chronic wounds that fail to properly heal, leading to the development of fibrosis (Liarte et al., 2020). TGF-β has also been shown to inhibit proliferation and induce the migratory phenotype acquired by wound-proximal keratinocytes (Sellheyer et al., 1993). Signal transduction is mediated by SMAD proteins, with Smad7 acting as a negative regulator of the TGF-β network (Schmierer and Hill, 2007). Evidence suggests that KGF levels are enhanced early in the healing response (Werner et al., 1992) whereas TGF-β displays a bimodal early/late distribution in the cutaneous wound (Liarte et al., 2020).

Longitudinal studies assessing wound healing during a tick bite should be coordinated by the scientific community because our studies were restricted to a single time point.

A critical component in the regeneration of the mammalian epidermis is the process of re-epithelialization. Keratinocytes proximal to the edge migrate towards the wound site, generating an adjacent gap that is replenished by proliferative cells (Park et al., 2017, Pastar et al., 2014).

Extracts from tick salivary glands have shown their capability to impede cellular growth *in vitro* (Hajnicka et al., 2011). Accordingly, we demonstrated that tick EVs led to a significant reduction in keratinocyte proliferation in both mouse and human cells. In mice, this observation was dependent on the role of DETCs, as DETC-deficient mice did not exhibit a decrease in proliferation of keratinocytes. Strikingly, the ability of ticks to impair epithelial proliferation was also observed in a quantitative-dependent manner. Increased number of ticks fed simultaneously at a given skin site resulted in a decrease of proliferative keratinocytes.

Wound healing in mice differs from humans (Zomer and Trentin, 2018). Mice utilize contraction mechanisms driven by the *panniculus carnosus* (Zomer and Trentin, 2018), whereas wound healing in humans relies on granulation tissue (Wang et al., 2024). Additionally, the Skint1-like gene whose mouse homolog has been implicated in the selection and seeding of DETCs to the epidermis is reportedly inactive in humans (Sutoh et al., 2018). Conversely, butyrophilins or butyrophilin-like molecules might enable the maturation and/or activation of skin-resident γδ T cells in both species (Boyden et al., 2008, Lewis et al., 2006, Ribot et al., 2021). Furthermore, epidermal γδ T cells from both mice and humans express the restricted Vδ1 TCR and skin-resident γδ T cells produce growth factors, which are important for epidermal homeostasis and wound healing (Bonneville et al., 2010, Holtmeier et al., 2001, Jameson et al., 2002, Ribot et al., 2021, Toulon et al., 2009). Therefore, it is possible that despite the technical limitations discussed in this article, tick EVs might affect γδ T cells and keratinocytes in both mammalian species.

## MATERIALS AND METHODS

### Reagents and resources

All primers, reagents, resources, and software used in this study, together with their manufacturer’s information and catalog numbers are listed in Supplementary Tables S1 and S14.

### Ticks

*I. scapularis* nymphs were obtained from two independent sources: (1) Dr. Ulrike Munderloh and Dr. Jonathan Oliver at the University of Minnesota; and the (2) tick rearing facility at Oklahoma State University. *A. americanum* and *D. variabilis* nymphs were obtained from the tick rearing facility at Oklahoma State University. Partially engorged *I. scapularis* adult ticks were obtained from Dr. Albert Mulenga and Dr. Adela Oliva Chavez at Texas A&M University. Upon arrival, ticks were maintained in a Percival I-30BLL incubator at 23°C with 85% relative humidity and a 12/10-hours light/dark photoperiod regimen.

### Mice

Experiments were performed on C57BL/6, FVB/N Jax, and FVB/N Tac mice. Breeding pairs were purchased from the Jackson Laboratory except FVB/N Tac mice, which were purchased from Taconic Biosciences. All mouse strains were bred at the University of Maryland School of Medicine, unless otherwise indicated. Male mice (7–9 weeks) were used for all experiments. All mouse experiments were approved by the Institutional Biosafety (IBC-00002247) and Animal Care and Use (IACUC numbers 0119012 and 1121014) committees at the University of Maryland School of Medicine and complied with the National Institutes of Health (NIH) guidelines (Office of Laboratory Animal Welfare [OLAW] assurance number A3200-01). *huLangerin-CreER;Rosa-stop-tdTomato;CX3CR1-GFP^+/−^;K14-H2B-Cerulean* mice used for intravital microscopy imaging were housed at Michigan State University as described elsewhere (Park et al., 2021) (IACUC number PROTO202300065). To activate DETCs, cytochalasin D (Sigma-Aldrich, C8273) was delivered topically as previously described (Park et al., 2021). Briefly, cytochalasin D was dissolved in a 25 mg/ml stock solution in dimethyl sulfoxide (DMSO), and later, the stock solution was diluted 100 times in 100% petroleum jelly (Vaseline; final concentration is 250 μg/ml). One hundred micrograms of the mixture of cytochalasin D and petroleum jelly were spread evenly on the skin once every 24 hours for 2 days. A mixture of 100% DMSO in petroleum jelly (1:100) was used as vehicle control.

### RNA interference

siRNAs and scRNAs for *vamp33* were designed as previously described (Oliva Chávez et al., 2021). Both siRNAs and scRNAs were synthesized according to the Silencer® SiRNA construction kit (Thermo Fisher Scientific). Primers are described in Supplementary Table S1.

Unfed nymphs were microinjected with 60-80 ng of siRNA or scRNA using a Nanoject III (Drummond Scientific Company). Ticks recovered overnight at 23°C with saturated humidity before being placed on respective mice.

### EV-depleted media

L15C300 medium was supplemented with 5% FBS (Millipore-Sigma), 5% tryptose phosphate broth (TPB) (BD), 0.1% lipoprotein concentrate (LPC) (MP Biomedicals), 0.25% sodium bicarbonate (Millipore-Sigma), and 25 mM HEPES (Millipore-Sigma). Media was cleared from EVs by ultracentrifugation at 100,000×g for 18 h at 4 °C in a LE-80 ultracentrifuge (Beckman Coulter) with a 60Ti rotor. EV-free media was then passed through a 0.22-μm Millipore Express® PLUS (Millipore-Sigma). The absence of EVs was confirmed by determining the particle size distribution with the NanoSight NS300 (Malvern Panalytical) for nanoparticle tracking analysis (NTA).

### Tick salivary gland culture

Salivary gland EVs were purified from *ex vivo* cultures that originated from partially engorged adult female ticks. Adult *I. scapularis* females were fed on New Zealand white rabbits for 5–6 days at either Dr. Albert Mulenga or Dr. Adela Oliva Chavez laboratories at Texas A&M University, as previously described (Oliva Chávez et al., 2021). Then, ticks were shipped to the University of Maryland School of Medicine. Partially-fed adult female ticks (90-120) were dissected 1–2 days post-removal. Briefly, midguts, Malpighian tubes, and other organs were removed. PBS was added to samples to avoid desiccation. Salivary glands were dissected and cultured in 24-well cell culture plates (Corning). 10 salivary glands from adult ticks were placed in each well, containing 500 μl of L15C300 EV-free medium supplemented with 1x penicillin/streptomycin (Corning) and 1x Amphotericin B (Gibco). Salivary glands were incubated for 24 h at 34 °C to allow EV secretion.

### EV purification

Medium collected from salivary gland cultures were cleared of any live cells by centrifugation at 300 × g for 10 minutes at 4 °C. Dead cells were removed by a second centrifugation at 2,000 × g for 10 minutes at 4 °C. The supernatant was collected, and apoptotic bodies were removed by a third centrifugation at 10,000 × g for 30 minutes at 10°C. The supernatant was filtered through a 0.22-μm Millipore syringe filter (Millipore-Sigma) to reduce the number of EVs >200 nm in size. EVs were pelleted by ultracentrifugation (100,000 × g) for 18 hours at 4 °C. Supernatant was discarded and EVs were resuspended in PBS. EV concentration and sizes were determined using the NanoSight 300 machine (Malvern Panalytical) with the software versions 2.0 or 3.0. The mean of the size generated in the reports was used to calculate the average size of the EVs in each sample. The concentration of proteins in tick EVs was determined using the BCA assay (Thermo Scientific), following the manufacturer’s procedure.

### Mouse capsule placement

Capsules made from the upper portion of a snap or screw top tube were adhered to the dorsal neck of each mouse to contain the ticks in one area. This technique is referred to as the capsule-feeding method and was adapted from a previous study (Schoeler et al., 1999). Briefly, capsule adhesive solution was made from 3 parts gum rosin (Sigma-Aldrich) and 1 part beeswax (FisherScience). Mice were anesthetized using isoflurane and shaved between the shoulder blades to the top of the cranium. Capsules were applied with the warmed adhesive and allowed to dry up for 24 hours prior to tick placement. Capsules were sealed with either a glued piece of mesh or a screw top after tick placement. Naïve groups consisted of capsule placement without ticks.

### Tick feeding experiments

Microinjected ticks were placed on mice using either the free-feeding or capsule-feeding method and allowed to feed for 3 days. On day 3, ticks were collected, weighed, and either placed in a humidified chamber for survival analysis or frozen at −80°C for RNA purification. To purify the mRNA, ticks were flash-frozen in liquid nitrogen and crushed with small plastic pestles. TRIzol® reagent (200 μl) was added to the crushed tick and RNA was purified using the PureLink™ RNA mini kit. cDNA was synthesized from 50 to 200 ηg (5–10 μl) of RNA using the Verso cDNA synthesis kit (Thermo scientific).

### Quantitative reverse transcription polymerase chain reaction (qRT-PCR)

qRT-PCR was performed to measure gene expression. qRT-PCR was performed with the CFX96 Touch Real-Time PCR Detection 233 System (Biorad). No template controls were included to verify the absence of primer-dimers formation and/or contamination. Reactions on each sample and controls were run in duplicate. Gene expression was determined by relative quantification normalized to tick *actin* using the primers listed in Supplementary Table S1.

### Flow cytometry of skin cell populations

*I. scapularis* nymphs fed on C57BL/6, FVB/N Jax, or FVB/N Tac male mice. On the third day of feeding, mice were euthanized with CO_2_. A 10- or 5-mm skin punch biopsy was taken while ticks were still attached. Skin samples from un-infested control mice were collected from matching locations. Single cell suspensions were prepared from each skin sample. Briefly, skin samples were cut into small pieces with sterile surgical scissors and placed into round-bottom tubes containing digestion buffer consisting of 90% RPMI-1640 (Quality Biological), 10% Liberase™ TL Research Grade (Roche), and 0.1% DNAse I (Millipore-Sigma). Digestions were carried out for 1 hour and 15 minutes at 37°C with constant shaking. Single cell suspensions were obtained by passing the digested tissues through a 40-μm cell strainer (Corning), homogenizing the tissue with a plunger and flushing cells with wash buffer consisting of PBS and 2 mM EDTA. Cells were centrifuged at 300 x g for 5 minutes at 4 °C, resuspended in 1 ml FACS buffer (PBS containing 1% BSA, 2 mM EDTA, and 0.05% NaN3) or FACS intracellular buffer (PBS containing 1% BSA and 0.05% NaN3). Cell suspensions were placed into a 96-well U-bottom plate and stained with respective antibody panels.

Live and dead cells were discriminated against using Zombie Violet Fixable Live Dead stain (BioLegend). Cells were washed with FACS buffer. Cells were then blocked with anti-FcR (CD16-CD32) (BioLegend 156603) and subsequently stained with the respective antibody panel for 15 minutes at 4°C and washed with FACS buffer. Whenever appropriate, anti-rat IgM was added to the cells, incubated for 15 minutes at 4°C, and washed twice with the FACS buffer.

Finally, cells were resuspended in 4% paraformaldehyde. For intracellular staining, cells were further processed following the instructions for the BioLegend’s FOXP3 Fix/Perm Buffer Set kit. Cells were measured with a LSRII flow cytometer (BD) at the Flow & Mass Cytometry Facility at the University of Maryland School of Medicine. Analysis was performed using the FlowJo software.

DETC populations in the murine skin were labeled with APC anti-CD45 (BioLegend 103111) or PE/Cyanine7 anti-CD45 (BioLegend 103114), FITC anti-CD3 (BioLegend 100203), BV60 anti-Vγ5 (BD 743241), APC anti-Thy1.2 (BioLegend 105312), and/or monoclonal antibody 17D1 (kindly provided by Dr. Adrian Hayday, King’s College London, and Dr. Robert Tigelaar, Yale University), and PE mouse anti-rat IgM (BD 553888). DETC costimulatory markers were measured with PE anti-JAML (BioLegend 128503), BV711 anti-CD100 (BD 745492), PE/Cyanine5 anti-CD44 (BioLegend 103010), APC/Cyanine7 anti-CD25 (BioLegend 102026), PerCP/Cyanine5.5 anti-CD69 (BioLegend 104522), and APC anti-CD314 (BioLegend 130212). Keratinocyte populations in the murine skin were labeled with BV711 anti-CD324 (BioLegend 118233), PE anti-CD200 (BioLegend 123807), PE/Cyanine5 anti-CD34 (BioLegend 119312), BV605 Sca1 (BioLegend 108133), and/or PE anti-CD49f (BioLegend 313612).

Keratinocyte proliferation was labeled with the Alexa Fluor 700 anti-Ki-67 (BioLegend 652420) in mice, and PE anti-Ki67 (BD 570922) in HaCaT cells. Smad7 was labeled with the anti-MADH7/SMAD7 polyclonal antibody (Abcam ab216428) and Alexa Fluor 405 goat anti-Rabbit IgG secondary antibody (Thermo Fischer Scientific A-31556).

### Enzyme-linked immunosorbent assay

To determine levels of KGF-1, 5 mm skin biopsies were placed in 200 µL of RPMI media tissue bath for one hour shaking at 32°C (150 revolutions per minute). KGF-1 levels in tissue bath supernatant were determined using the human R&D Systems KGF/FGF-7 Quantikine ELISA kit according to manufacturer instructions (Supplementary Table S14). To determine levels of TGF-β, 5 mm skin biopsies were homogenized in lysis buffer containing 1X RIPA buffer (catalogue number 20-188, Millipore) with 1X protease-phosphatase inhibitor cocktail (catalogue number 78420, Thermo Scientific). TGF-β levels in tissue supernatant were determined using the R&D Systems TGF-β 1 Quantikine ELISA kit according to manufacturer instructions. Total protein in samples was determined using the Pierce BCA Protein Assay Kit (catalogue number 23227 Thermo Scientific). Concentration of KGF-1 and TGF-β were normalized to the total protein in a sample.

### Intravital microscopy

Epidermal intravital imaging studies were done in collaboration with Dr. Sangbum Park at Michigan State University. All *in vivo* imaging and analysis were performed, as described previously (Park et al., 2021). Simultaneous visualization of Langerhans cells, DETCs and epithelial cells was achieved by utilizing the *huLangerin-CreER;Rosa-stop-tdTomato;CX3CR1-GFP^+/−^;K14-H2B-Cerulean* mice.

### Epidermal single-cell isolation, scRNA-seq library preparation and sequencing

*I. scapularis* nymphs were microinjected with *vamp33* si or *vamp33* sc and fed on FVB/N Jax or FVB/N Tac mice. On the third day of feeding, mice were euthanized with CO_2_. Partially fed ticks were removed and the sites where ticks bit were shaved followed by an application of a light layer of Nair depilatory lotion. A total of three 5-mm skin punch biopsies were obtained from the dorsal neck for each mouse. 5-mm skin punch biopsies were obtained from the same physiological site of naïve mice. Skin samples were incubated in dispase solution (4 U/mL dispase, 5 mM MgCl2, and 0.4 mM CaCl2 in PBS) for 2.5 hours at 37°C with constant shaking/stirring. Epidermal sheets were separated from the dermal layer using forceps. Epidermal sheets were then incubated in a digestion solution (2.5 mg/mL collagenase D and 0.2 mg/mL DNase in RPMI Medium) for 1 hour at 37°C with constant shaking/stirring.

Cells were resuspended using a wide-bore pipette tip and three samples per treatment per mouse were combined. Samples were passed through a 40 µM cell strainer and washed with RPMI +10% FBS. Cells were counted using the Countess II FL Automated Cell Counter, stained with 5 μl of 7-AAD per million cells, and incubated in the dark for 10 minutes at 4°C. Samples were then sorted at the CIBR Flow Cytometry Core Facility at the University of Maryland School of Medicine. Cells were sorted into a PBS in the absence of calcium and magnesium + 10% FBS collection buffer. They were then transported on ice to the Institute of Genome Sciences at the University of Maryland School of Medicine for library preparation and sequencing. Single cell libraries were generated with the 3’ NextGEM v3.1 kit targeting 3800-5000 cells. Libraries were sequenced with a NovaSeq 6000, S2 flowcell targeting 375M read pairs per sample.

### Bioinformatics

All scRNA-seq reads were processed and mapped to the mouse mm10 reference genome using 10X Genomics’ Cell Ranger software. Approximately 20,640 total cells were profiled with 88,027 mean reads per cell across all conditions. A count matrix (gene-by-cell) generated by cell ranger count for each library was then aggregated into a single count matrix. Expression matrices were generated using the Bioconductor packages scater (v1.22.0) (Lun et al., 2016b) and scran (v1.22.1) (Lun et al., 2016a). Cells with less than 2,500 or greater than 60,000 UMIs were removed after calculating cell metrics using scater (v1.22.0). DoubletFinder (v2.0.1) (McGinnis et al., 2019) was applied removing 1,364 cells, which yielded a total of 10,715 cells. The remaining transcriptomes were normalized by first calculating size factors via the scran functions quickCluster and computeSumFactors. Then, we computed normalized counts for each cell with logNormCounts function in scran (v1.22.1).

For downstream analysis, highly variable genes were selected using getTopHVGs before performing the Principal Component Analysis (PCA) and the tSNE projection. Clustering was conducted using kmeans function based on the calculated tSNE. Differential gene expression between clusters was calculated using find Markers function. Only identified epidermal cells of interest (Keratinocytes, T cells, and APCs) were further analyzed, resulting in a total of 5,172 cells with a median UMI count of 13,910 per cell. For pseudotime analysis, the Bioconductor matrix was imported into slingshot (v2.2.1) (Street et al., 2018). To compare the T cell receptor delta variable 4 (*Trdv4*) expression, normalized counts were used for visualization by the violin plot. The permutation test was applied to calculate the significance of the difference in the mean expression between two groups. A list of differentially expressed keratinocyte genes between treatments was generated by MAST (v1.24.0) (Finak et al., 2015) with significance testing under the Hurdle model for downstream analysis by the IPA.

### Gene set enrichment analysis

Gene set enrichment analysis was performed using DAVID, version 2021. Default DAVID parameters were employed and included the following categories for the enrichment analysis: GOTERM_BP_DIRECT, GOTERM_CC_DIRECT and GOTERM_MF_DIRECT (from Gene_Ontology), KEGG_PATHWAY (from Pathways) and INTERPRO (from Protein_Domains). *p* value and FDR< 0.05 were set as a threshold.

### Ingenuity pathway analysis

Differentially expressed keratinocyte genes from the following samples were analyzed in the IPA as independent datasets: 1) FVB-Tac Naïve versus FVB-Jax Naïve 2) FVB-Jax si*V33* versus FVB-Jax sc*V33* and 3) FVB-Tac si*V33* versus FVB-Tac sc*V33*. Genes were considered differentially expressed if the *p* value and FDR were < 0.05. Dataset input criteria for the IPA included expression, *p* value, log ratio, FDR, and Ensemble ID codes. All datasets were examined for canonical pathway and upstream regulator analysis. FVB-Tac Naïve versus FVB-Jax Naïve dataset had 591 IDs, including 589 mapped and 2 unmapped IDs. FVB-Jax si*V33* versus FVB-Jax sc*V33* dataset had 1207 IDs, including 1204 mapped and 3 unmapped IDs. FVB-Tac si*V33* versus FVB-Tac sc*V33* had 732 IDs, including 728 mapped and 4 unmapped IDs. The IPA proprietary algorithm segments the network map between molecules into multiple networks and assigns scores for each network as described previously (Calvano et al., 2005). For the canonical pathway analysis, −log (P-value) >2 was taken as threshold and for the upstream regulator analysis, the *p* value of overlap <0.05 was set as the threshold. A positive *Z*-score was defined as the predicted activation, and a negative *Z*-score was defined as the predicted inhibition.

### Epithelial-to-mesenchymal transition signature score

The “AddModuleScore” (Seurat, version 4.3.0) function was used to generate an epithelial-to-mesenchymal transition score across all keratinocytes based on 102 genes reported (Guo et al., 2024).

### Cell culture

Human keratinocyte HaCaT cells were grown in T75 flasks (CytoOne CC7682-4875) and maintained in Dulbecco’s Modified Eagle Medium (DMEM) (Gibco 10564011) supplemented with 10% fetal bovine serum (FBS) (Millipore-Sigma F0926-500ML) with 1% penicillin-streptomycin (Gibco, 15140122).

### Scratch wound assay

HaCaT cells were seeded into 24-well plates at a density of 1x10^6^ cells/mL and cultured to confluency. Scratch wounds were inflicted using the SPLScar 24-well scratcher (SPL Life Sciences 201924). For gap closure, media was replaced with or without *I. scapularis* EVs at three concentrations (10^2^/mL, 10^4^/mL, or 10^6^/mL) and the gap area was imaged at 0-, 12-, and 24-hours post-scratch with the AMG EVOS fluorescence microscope and quantified with ImageJ software. For proliferation studies, media was replaced with or without *I. scapularis* EVs at 10^2^/mL, or 100 µg/mL mitomycin C as a proliferative control. At 24 hours post-scratch, cells were collected for Ki67^+^ staining.

### Statistical analysis

Statistical significance was assessed as follows: percent tick attachment was calculated by the Fisher’s exact test, tick weight by the *t* test or the Mann Whitney test, and survival curve by the Log-rank (Mantel-Cox) test. One-way ANOVA followed by Tukey’s *post hoc* test for multiple comparisons was also used. Kruskal-Wallis ANOVA was implemented if the dataset failed normality of residuals or displayed heterogeneity of variance. We used GraphPad PRISM® (version 9.1.0) for all statistical analyses. Outliers were detected by a GraphPad Quickcals program (https://www.graphpad.com/quickcalcs/Grubbs1.cfm). *p* values of < 0.05 were considered statistically significant.

## DATA AVAILABILITY

All scRNA sequences are deposited into the NCBI Sequence Read Archive under the BioProject accession PRJNA905677. R codes for scRNA sequencing datasets were adapted from https://bioconductor.org/books/3.16/OSCA/ and specified R package vignettes. Tokens can be made available upon request. Further information and request for resources and reagents should be directed to and will be honored by the corresponding author: Joao HF Pedra

## CONFLICT OF INTEREST

None

## ACKNOWLEDGEMENTS

We acknowledge members of the Pedra laboratory for providing insightful discussions. We thank the rearing facility at Oklahoma State University for providing *I. scapularis, A. americanum,* and *D. variabilis* ticks; Xiaoxuan Fan, Bryan Hahn, Regina Harley, and Sean McGill (University of Maryland School of Medicine) for flow cytometry and sorting assistance; Adrian Hayday (King’s College London) and Robert Tigelaar (Yale University) for the monoclonal antibody 17D1; the Maryland Genomics Core at the Institute for Genome Sciences, University of Maryland School of Medicine for the services provided in next generation sequencing; the University of Maryland Greenebaum Comprehensive Cancer Center Flow Cytometry Shared Service core facility and the Flow & Mass Cytometry Facility at the University of Maryland School of Medicine for flow cytometry services; Cristiana Cairo, Nevil Singh, Nicholas Carbonetti (University of Maryland School of Medicine) and Jere McBride (University of Texas Medical Branch) for insightful advice. This work was supported by grants from the NIH to F31AI152215 (AJO), F31AI167471 (LRB), R01AI134696 (JHFP), R01AI116523 (JHFP), P01AI138949 (JHFP), T32AI162579 (HJL-Y), T32AI095190 (DMR), R01AR083086 (SP), Hatch-Multistate Project to TEX0-1-7714 (ASOC), the Diseases of Agriculture Animals program (A1221) from the United States Department of Agriculture, National Institute of Food and Agriculture (USDA-NIFA) award #2022-67015-42166 (ASOC), the Knipling-Bushland-Swahrf fellowship from the Department of Entomology at Texas A&M University (BL-G). The content is solely the responsibility of the authors and does not represent the official views of the NIH, the Department of Health and Human Services, the USDA-NIFA or the United States government.

## AUTHOR CONTRIBUTIONS

Conceptualization: LM, LMV, JHFP; Data Curation: LM, LMV, HNB, HJL-Y, AR; Formal Analysis: LM, LMV, HNB, HJL-Y, AR, CRF; Funding Acquisition: JHFP; Investigation: LM, LMV, CRF, AJO, ADS-S, DMR, MTM, SS, LRB, NS, FECP, AW-C; Methodology: LM, LMV, JHFP; Project Administration: LM, LMV, JHFP; Resources: EBC, JMJ, T-TN, BL-G, JDO, UGM, ASOC, AM, SP, DS, JHFP; Software: HNB, HJL-Y, AR, ADS-S, SP, DS; Supervision: SP, DS, JHFP; Validation: LM, LMV; Visualization: LM, LMV, LRB; Writing – Original Draft Preparation: LM, LMV, JHFP; Writing – Reviewing and Editing: LMV, JHFP.

## Notes

### Competing Interest Statement

The authors have declared no competing interest.

### Summary of Updates

We: (1) clarified our previously established model for the reduction of tick extracellular vesicles (EVs) upon Vamp33 siRNA silencing; (2) showed a diminished capacity of gap closure upon tick EV stimulation in a scratch wound assay (Figure 6a-c); (3) provided additional evidence of an EV-mediated reduction in keratinocyte proliferation (Figure 6d); (4) enhanced our computational analysis of the unidentified subpopulation to reveal a prominent epithelial-mesenchymal transition gene signature (Figure 4f; Supplementary Figure S8 and Supplementary Table S10); (5) implemented more judicious language throughout the manuscript to temper overstated conclusions; (6) expanded the rationale for the tick EV and mouse DETC-deficient models used; and (7) used the discussion section to explain the constraints associated with the tick-epidermal system.

