## Supplementary Figure Legends for "Tick extracellular vesicles alter keratinocyte function in the skin epidermis"

**Supplementary Figure S1: DETC flow cytometry gating strategy.** 5 mm skin punch biopsies were obtained from the bite of ticks and compared to the naïve skin followed by flow cytometry analysis. Representative flow cytometry plots were gated for **(a)** DETCs (Vy5^+^) and **(b)** DETC

co-receptors (JAML^+^, NKG2D^+^, CD25^+^, CD69^+^, CD100^+^ or CD44^+^).

**Supplementary Figure S2: ScRNA-seq data filtration.** Composite datasets of FVB-Jax and FVB-Tac samples included 20,640 cells **(a)** before filtration by scran (R package). **(b)** tSNE plot of fixed threshold filtration, set to 2500-60,000 UMIs. **(c)** Doublet finder (R package) of dataset. tSNE was colored by the doublet score. **(d)** tSNE plot after fixed threshold filtration and doublet finder analysis.

**Supplementary Figure S3: Expression of keratinocyte-specific markers. (a)** Graphical illustration of keratinocyte stratified layers with select marker genes. tSNE of keratinocyte clusters depicting gene expression of **(b)** *Krt14*, **(c)** *Krt5*, **(d)** *Krt1*, **(e)** *Krt10* and **(f)** *Ivl*.

**Supplementary Figure S4: Expression of hair follicle-specific markers. (a)** Graphical illustration of hair follicle microanatomy with select marker genes. tSNE of keratinocyte clusters depicting gene expression of **(b)** *Shh*, **(c)** *Krt75*, **(d)** *Lgr5*, **(e)** *Mgst1* and **(f)** *Krt79*.

**Supplementary Figure S5: Epidermal cell type characterization.** Cluster frequency of keratinocytes, antigen presenting and T cells in **(a)** FVB-Jax and **(b)** FVB-Tac mice in the presence or absence of *I. scapularis* nymphs microinjected with *scV33* or *siV33*. **(c)** Violin plot displaying the expression of the TCR-Vδ1 gene, *Trdv4*, in the epidermal T cell cluster of naïve FVB-Jax and FVB-Tac mice. Significance shown as **p*<0.05 based on a permutation test using R statistical packages.

**Supplementary Figure S6: Individual tSNE plots of keratinocyte clusters**. Subclustering analysis of keratinocytes across samples: **(a)** FVB-Jax, **(b)** FVB-Jax *scV33*, **(c)** FVB-Jax *siV33*, **(d)** FVB-Tac, **(e)** FVB-Tac *scV33*, and **(f)** FVB-Tac *siV33*.

**Supplementary Figure S7: Keratinocyte-specific markers along pseudotime trajectory. (a)** *Krt14*, **(b)** *Krt1*, and **(c)** *Ivl* gene expression along pseudotime values (x axis) for naïve, *scV33*-, or *siV33*-tick bites on FVB-Jax or FVB-Tac mice. Cells colored by clusters from keratinocyte tSNE plot (as shown in Figure **4a**) ordered across the pseudotime (x-axis).

**Supplementary Figure S8: Epithelial-to-mesenchymal transition score across all scRNAseq clusters.** The “AddModuleScore” function was used to generate an epithelial-to-mesenchymal transition score (EMT) across all clusters after final filtration (Supplementary Figure S2d). Epithelial-to-mesenchymal transition score was based on 102 epithelial-to-mesenchymal genes (Guo et al., 2024). Selected genes were reported as indicative of keratinocytes in intermediary or mesenchymal transitioning states.

**Supplementary Figure S9: Flow cytometry gating strategy in keratinocytes.** 5 mm punch biopsies were obtained from the bite site of ticks or naïve skin and processed for flow cytometry. Representative flow cytometry plots were gated for **(a)** EpCAM^+^ keratinocytes, **(b)** Ki67^+^, **(c)** Anti-rabbit IgG^+^ for PI3K and Smad7, and **(d)** p-PI3K^+^ keratinocytes.
